# Sequestration of TDP-43^216-414^ aggregates by cytoplasmic expression of the proSAAS chaperone

**DOI:** 10.1101/2020.04.15.039578

**Authors:** Juan R. Peinado, Kriti Chaplot, Timothy S. Jarvela, Edward M. Barbieri, James Shorter, Iris Lindberg

## Abstract

As neurons age, protein homeostasis becomes less efficient, resulting in misfolding and aggregation. Chaperone proteins perform vital functions in the maintenance of cellular proteostasis, and chaperone-based therapies that promote sequestration of toxic aggregates may prove useful in blocking the development of neurodegenerative disease. We previously demonstrated that proSAAS, a small secreted neuronal protein, exhibits potent chaperone activity against protein aggregation *in vitro*, and blocks the cytotoxic effects of amyloid and alpha synuclein oligomers in cell culture models. We now examine whether cytoplasmic expression of proSAAS results in interaction with protein aggregates in this cellular compartment. We used site-directed mutagenesis, confocal microscopy, *in vitro* aggregation assays, and functional assays to investigate the interaction of proSAAS with TDP-43 and other known aggregating proteins. We report that expression of proSAAS within the cytoplasm generates dense, membrane-less 2 μm proSAAS spheres which progressively fuse to form larger spheres, suggesting liquid droplet-like properties. ProSAAS spheres selectively accumulate a C-terminally truncated fluorescently-tagged form of TDP-43^216-414^, initiating its cellular redistribution by sequestration within the sphere core; these TDP-43^216-414^ -containing spheres also exhibit dynamic fusion. Removal of either the predicted α-helix (37-70) composed of hydrophobic and charged amino acids or the stretch of amino acids encompassing the conserved hydrophobic region and the positively charged furin site (163-189) inhibits the ability of proSAAS both to form spheres and to encapsulate TDP-43 aggregates. As a functional output, we demonstrate that proSAAS expression results in cytoprotection against full-length TDP-43 toxicity in yeast. In summary, the normally secreted neuronal chaperone proSAAS, when expressed in the cytoplasm unexpectedly phase-separates to form spherical liquid-like condensates that undergo dynamic fusion. We conclude that cyto-proSAAS acts as a functional holdase for cytoplasmic TDP-43^216-414^ molecules via this phase-separation property, representing a cytoprotectant whose unusual biochemical properties can potentially be exploited in the design of therapeutic molecules.

## INTRODUCTION

In recent years it has become increasingly apparent that many, if not all neurodegenerative diseases involve the aggregation of key proteins, including Alzheimer’s, Parkinson’s, Huntington’s diseases as well as amyotrophic lateral sclerosis (ALS). During disease progression, misfolded proteins, many containing intrinsically disordered domains, initiate a proteostatic cascade in which proteins are destabilized and precipitate, contributing to neurotoxic events that culminate in neuronal death ^1, 2^. Examples of cytoplasmic aggregates involved in neuropathology include aggregated alpha-synuclein (α-Syn), the major component of the cellular hallmark of Parkinson’s disease, Lewy bodies; and the RNA-binding protein TDP-43, which normally shuttles between the cytoplasm and nucleus, but is mislocalized to cytoplasmic aggregates in more than 95% of patients with ALS, and in 45% of those with frontotemporal dementia (^3^; ^4^; reviewed in ^5^). TDP-43 is post-translationally modified by phosphorylation and ubiquitination as well as by removal of its aggregation-prone carboxy-terminal prion-like domain, and all of these modifications are strongly implicated in disease pathology ^4^. These abnormal proteins undergo liquid-liquid phase separation (LLPS) to form liquid condensates *en route* to the formation of solid phase aggregates (e.g. ^6^); LLPS is thought to be important to proteostatic control during cellular stress ^7^.

Increasing evidence points to important roles for cellular and secreted chaperones in the formation and deposition of abnormal protein aggregates in neurodegenerative disease [reviewed in ^2, 8-10^]. Chaperone proteins may act to sequester misfolded proteins; refold unfolded proteins; and cooperate with degradative machinery, either directly or indirectly, to facilitate the degradation of misfolded species and/or enhance disaggregation (see above reviews). Overexpression of cytoplasmic chaperones mitigates aggregate toxicity ^2, 11, 12^, whereas overexpression of the constitutively-secreted secretory chaperones clusterin and α2-macroglobulin is neuroprotective in Neuro2A cells ^13^; reviewed in ^14^.

Despite the enhanced susceptibility of neurons to proteostatic disease, few neuron-specific chaperones have been identified in brain tissue. We have previously reported that the neuronally-expressed secretory chaperone proSAAS potently blocks the fibrillation of neurodegenerative disease-related proteins such as β-amyloid and α-Syn and is cytoprotective against oligomers of these two proteins ^15, 16^. ProSAAS immunoreactivity colocalizes with inclusion bodies in a variety of neurodegenerative diseases ^15-18^, supporting a role for proSAAS in disease control. Interestingly, ten independent proteomics studies have identified proSAAS as a differentially expressed biomarker in cerebrospinal fluid obtained from dementia patients ^19-28^. Recent transcriptomics studies of Alzheimer’s brain tissue show that proSAAS expression increases during disease progression ^29^; and other studies indicate that proSAAS synthesis is highly increased in a hippocampal model of homeostatic scaling ^30^.

These data implicating proSAAS in neurodegenerative disease have prompted us to investigate a potential role for the proSAAS chaperone in handling misfolded proteins involved in neurodegeneration-related processes in living cells. Unexpectedly, we found that cytoplasmic expression of proSAAS results in the formation of unique spherical structures that exhibit high affinity for TDP-43^216-414^, with beneficial consequences.

## RESULTS

To explore the role of proSAAS as a chaperone in neurodegenerative diseases, we performed cell culture-based co-expression studies to identify potential client proteins that are prone to aggregation. We expressed secreted FLAG-tagged proSAAS (“sec-proSAAS”, **Figure 1A**), which naturally localizes to the secretory pathway of HEK and Neuro2A cells, but since many aggregating proteins involved in neurodegeneration are cytoplasmic, we also expressed a FLAG-tagged proSAAS construct lacking a signal peptide (“cyto-proSAAS”; **Figure 1A**).

**Figure 1.**
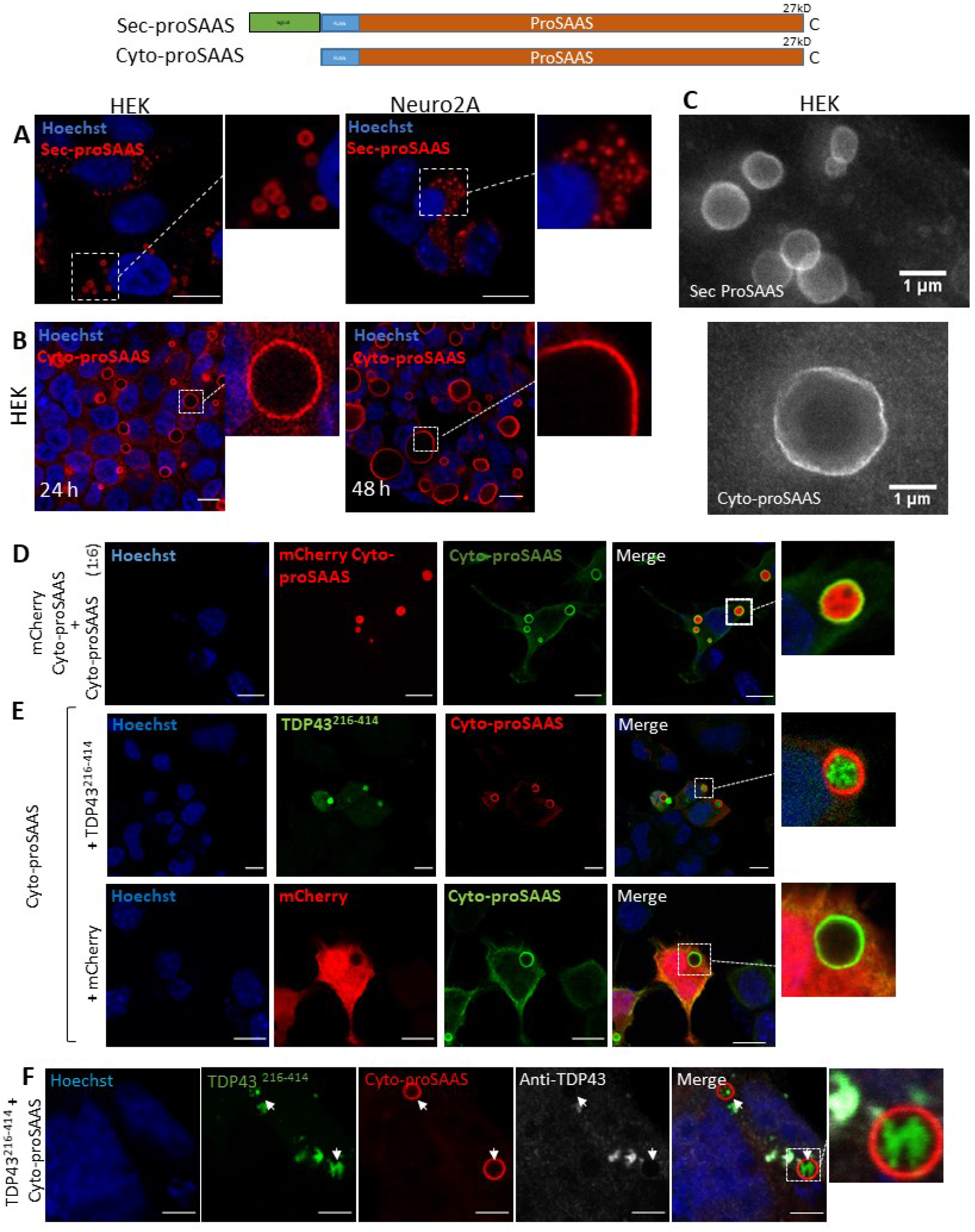
Cytoplasmic expression of proSAAS leads to the formation of dense antibody-impenetrable spheres that specifically encapsulate TDP-43^216-414^ aggregates. Constructs used are shown above the figure, with the FLAG tag shown in blue and the signal peptide in green. **Panel A:** Immunostaining of secretory proSAAS (sec-proSAAS) expressed for 48 h (*red*) reveals the appearance of spheres in HEK cells (**left**). Neuro2A cells rarely contain these spheres **(right)**. **Panel B:** Representative confocal images of HEK cells transfected with cyto-proSAAS cDNA. 48 h after transfection the spheres are larger in size than at 24h (**left vs right sides**). **Panel C:** Representative STED images of flag-immunoreactive spheres formed by overexpressed secretory proSAAS (*top*) and cyto-proSAAS (*bottom*) in HEK cells. **Panel D:** Co-transfection of small amounts of mCherry-cyto-proSAAS with a large (five-fold) excess of cyto-proSAAS (*green*) results in dense fluorescent mCherry-tagged proSAAS spheres. **Panel E (top row):** TDP-43^216-414^ aggregates are found inside cyto-proSAAS spheres (*red*) when both are co-expressed in Neuro2A cells. **Panel E (bottom row)** shows the characteristic cytosolic fluorescence of the control protein mCherry (*red*), which is not accumulated within the cyto-proSAAS spheres *(green*) when both are co-expressed in Neuro2A cells. **Panel F:** Lack of immunostaining of Neuro2A cells with GFP-TDP-43 antibody (C-terminal; in *white*) after co-expression of cyto-proSAAS (*red*) and TDP-43^216-414^ (*green*), shows that this antibody cannot penetrate inside the sphere, supporting core inaccessibility. **Arrow** indicates fluorescent, but immunologically undetectable TDP-43^216-414^ aggregates inside spheres. Images taken at 40X, 24 h after transfection. Scale bar, 10 μm. Magnified inserts correspond to 10 μm.

### Cytoplasmic expression of proSAAS results in the formation of dense antibody-impenetrable spherical structures that specifically encapsulate GFP-TDP-43^216-414^ aggregates

Following transfection of sec-proSAAS into Neuro2A cells and staining with FLAG antibody, we observed the expected punctate localization, likely within secretory vesicles (**Figure 1A)**. However, in sec-proSAAS-expressing HEK cells, proSAAS FLAG staining presented as 1 µm large spherical structures (**Figure 1A)**. Expression of the cyto-proSAAS construct in both HEK (**Figure 1B**) and Neuro2A cells (**Figure S1A**) resulted in the appearance of large FLAG-immunoreactive spherical structures in the cytosol. In Neuro2A cells, these spheres exhibited an average size of 2 ± 1 µm. Similar results were found in HEK cells at 24 h after transfection, though the size of the spheres was greater, with an average diameter of 4 ± 2 µm. This pattern of expression remained constant at 48 h in Neuro2A cells (**Figure S1A**), but in HEK cells the size of the spheres increased at this timepoint, suggesting increased proSAAS recruitment (**Figure 1B**). ProSAAS spheres were easily observable at 40x using phase-contrast microscopy (**Figure S1B**). Western blotting of cyto-proSAAS-transfected HEK cells revealed the expected band of approximately 30 kDa, corresponding to full-length proSAAS (**Figure S1B**). Transfected hippocampal neurons also exhibited cyto-proSAAS spheres (**Figure S1C**), suggesting that proSAAS sphere formation is not restricted to propagating cell lines.

We then performed STED imaging of sec-proSAAS sphere morphology, which confirmed the presence of 1 µm-sized spheres stained with FLAG antibody within the secretory pathway (**Figure 1C**). Similar experiments performed with cyto-proSAAS revealed the presence of an undulated and wavy surface on 2-4 µm large spheres (**Figure 1C**). Taken together, these data show that the proSAAS chaperone is not diffusely expressed either within the secretory pathway or the cytoplasm, but instead strongly self-associates, condensing into distinct spheres.

To investigate how proSAAS forms these spherical assemblies, we attached an amino-terminal mCherry tag to the cyto-proSAAS construct (“mCherry-cyto-proSAAS”) and transfected this construct into Neuro2A cells. Surprisingly, rather than forming spheres, mCherry-cyto-proSAAS formed aggregates dispersed throughout the cytosol, indicating possible misfolding (**Figure S1D**). Reducing the concentration of mCherry-cyto-proSAAS cDNA by dilution with a 6-fold excess of non-mCherry-tagged cyto-proSAAS cDNA resulted in the formation of dense cyto-proSAAS spheres which were uniformly labelled with mCherry fluorescence (**Figure 1D**). We surmise that excess cyto-proSAAS interacts with mCherry-cyto-proSAAS, enabling sphere formation.

To identify potential aggregating client proteins for the cyto-proSAAS chaperone, we expressed cyto-proSAAS together with cDNAs encoding various GFP-tagged cytoplasmic proteins implicated in neurodegenerative diseases and known to aggregate intracellularly, including synuclein, tau, huntingtin polyQ repeats (HTT-Q74) and TDP-43. GFP-tagged synuclein and tau, and their respective mutants synuclein A53T and TauE14, exhibited uniform cytoplasmic distribution, while HA-tagged and GFP-tagged HTT-Q74, and the TDP-43 C-terminal construct GFP-TDP-43^216-414^ formed dispersed cytoplasmic inclusions when expressed alone in HEK cells (**Figure S1E**). While the non-aggregating forms of synuclein and tau proteins failed to interact with cyto-proSAAS spheres, the distribution of GFP-TDP-43^216-414^ aggregates was profoundly altered by co-expression of cyto-proSAAS, becoming localized predominantly within the proSAAS sphere cores. This occurred both in Neuro2A (**Figure 1E, upper row**) and HEK cells (**Figure S2B)**. In contrast, neither HA-tagged HTT-Q74, nor GFP**-**HTT-Q74 aggregates were incorporated into spheres (**Figure S2C, S2D**), nor was mCherry incorporated (**Figure 1E, lower row; S2E)**. Using TDP-43 antibody staining, cytoplasmic GFP-TDP-43^216-414^ located outside cyto-proSAAS spheres, but not encapsulated GFP-TDP-43^216-414^ could be visualized, supporting the idea that cyto-proSAAS spheres are antibody-impenetrable (**Figure 1F**). These data indicate that the C-terminal portion of TDP43 exhibits a great affinity for the interior of the proSAAS sphere.

We then used transmission electron microscopy to gain a better structural understanding of proSAAS-TDP-43^216-414^ sphere structure (**Figure 2A, top**). When transfected alone, GFP-TDP-43^216-414^ accumulated in small amorphous structures of 100 to 500 nm (**Figure 2A, bottom)**. Cells transfected with both cyto-proSAAS and GFP-TDP-43^216-414^ cDNA showed electron-dense spheres of the expected size, in agreement with immunofluorescence experiments (**Figure 2B)**. The uniform electron density of these spheres did not reflect the polydispersity of internal GFP-TDP-43^216-414^ aggregates seen in fluorescent images. Untransfected cells contained neither the small amorphous structures nor the spheres (data not shown). These electron density data indicate that cyto-proSAAS spheres are not hollow but encapsulate material. This conclusion is supported by a 3D-rendered image which shows immunoreactive proSAAS surrounding GFP-TDP-43^216-414^ (**Figure 2C**).

**Figure 2.**
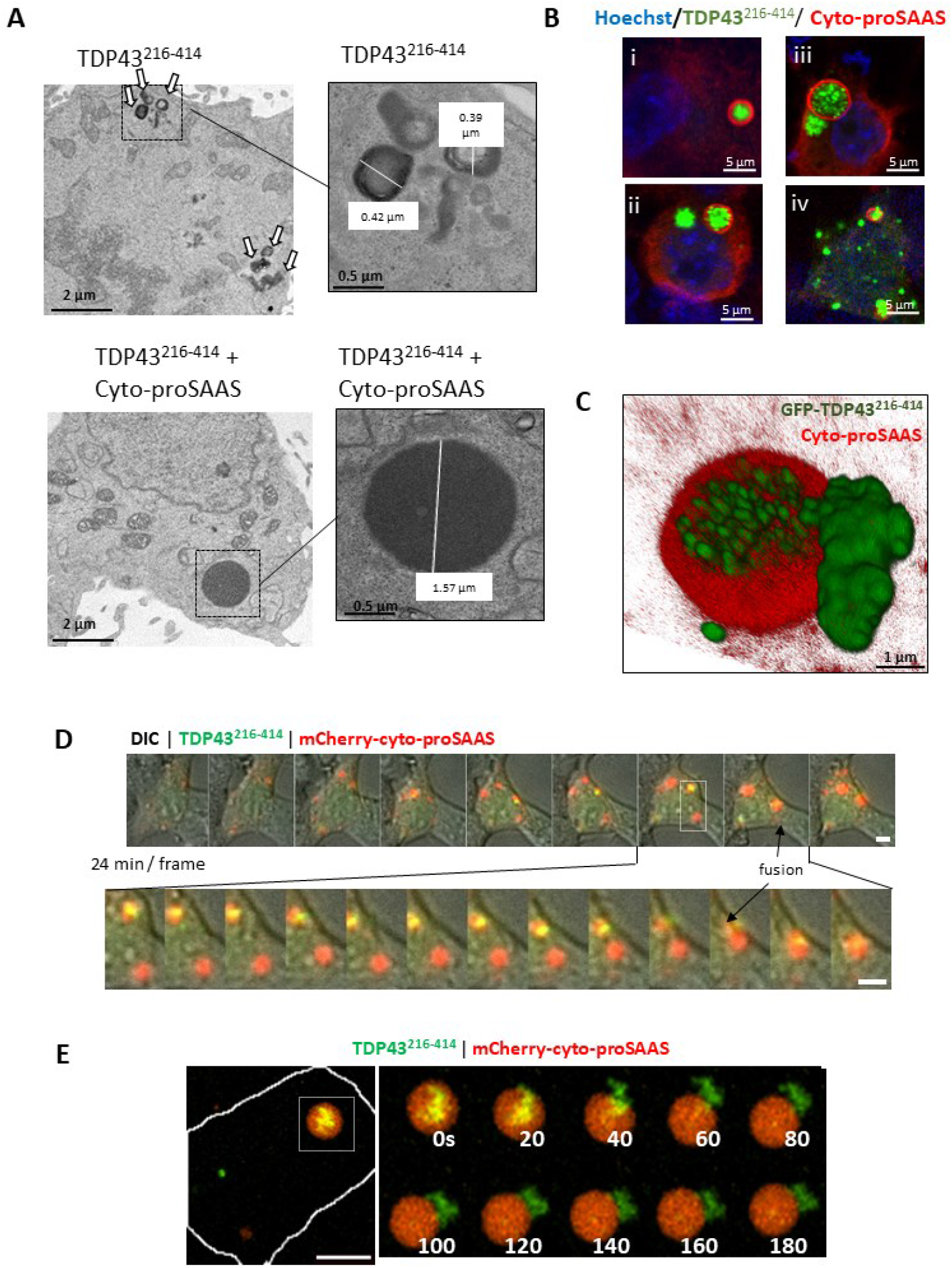
Cyto-proSAAS spheres are electron-dense and display simultaneous dynamic fusion and encapsulation of GFP-TDP-43^216-414^ aggregates. **Panel A (top)** EM images of HEK cells transfected only with GFP-TDP-43^216-414^. The characteristic GFP-TDP-43^216-414^ aggregates are indicated with arrows, and a magnified image is shown at right. **Panel A (bottom):** Electron microscopy images of HEK cells expressing GFP-TDP-43^216-414^ and cyto-proSAAS, showing the characteristic dense spheres. Scale bars of 10 mm are indicated in the images. *Red*, cyto-proSAAS, *green*, GFP-TDP-43^**216-414**^. **Panel B**. Confocal images of HEK cells co-transfected with GFP-TDP-43^216-414^ and cyto-proSAAS cDNAs showing differing TDP-43 morphologies. **Panel C**. 3D visualization of the spherical cyto-proSAAS-TDP-43^**216-414**^ assembly. **Panels D and E:** Dynamic widefield microscopy of Cherry-cyto-proSAAS (*red*) and GFP-TDP-43^216-414^ (*green*, with DIC) beginning 18h post transfection, taken at 2 min intervals for 5h. See also ***Supplemental Movies 1 and 2***. **Panel D**: Images, taken at 24 min intervals (*top*), depict the generation of multiple spheres, some containing GFP-TDP-43^216-414^ aggregates which condense and fuse into larger spheres. A subset of images (*box*) is shown below at 2 min intervals, which captures two large spheres condensing into one. **Panel E**: Rarely, internal GFP-TDP-43^216-414^ was observed to rapidly and wholly exit from the center of cyto-proSAAS spheres. Max projections are shown, but a 3D analysis reveals GFP-TDP-43^216-424^ to be contained entirely within the sphere at the beginning of the time course. Cyto-proSAAS, GFP-TDP-43^216-414^ and Cherry-cyto-proSAAS were co-transfected in a ratio 1:1:0.2, as indicated in *Methods*. Scale bar, 10 μm. Magnified inserts correspond to 10 μm.

### Dynamic fusion of liquid-like cyto-proSAAS spheres containing GFP-TDP-43^216-414^ aggregates indicates properties consistent with phase separation

To view the dynamics of cyto-proSAAS sphere formation, we performed widefield microscopy beginning 18 h after triple transfection with cDNAs encoding cyto-proSAAS, GFP-TDP-43^216-414^, and mCherry-cyto-proSAAS (in the weight ratio 1:1: 0.2; see *Materials and Methods*) of HEK cells. Cyto-proSAAS spheres first formed as several smaller spheres before fusing into only one or a few large spheres per cell, demonstrating liquid-liquid phase separation-like ability (**Figure 2D; see also Supplemental Movies 1, 2**). Small spheres also took up GFP-TDP-43^216-414^ aggregates while continuing to fuse further. Occasionally, but only rarely, we observed full extrusion of GFP-TDP-43^216-414^ aggregates from the center of cyto-proSAAS spheres (**Figure 2E**). When this occurred, the entirety of the GFP-TDP-43^216-414^ content was removed over the course of 40 seconds, showing that encapsulation is a reversible process. However, since most cellular GFP-TDP-43^216-414^ is encapsulated, extrusion is likely energetically less probable than uptake.

### Cyto-proSAAS spheres do not contain markers corresponding to known organelles

Cyto-proSAAS spheres appear to represent an unusual cytoplasmic structure. To gain information as to whether cyto-proSAAS spheres possess markers corresponding to known organelles, or fuse with known organelles, we stained sphere surfaces with antisera directed against various cellular markers characteristic of different organelles. LC3-mCherry, an autophagosomal marker, did not colocalize or become sequestered by cyto-proSAAS, indicating that cyto-proSAAS sphere formation does not pertain to autophagic degradation (**Figure S3A**). Markers of lysosomes (Lamp-1) or Golgi (giantin) (**Figure S3B-S3C**) also failed to stain cyto-proSAAS spheres. Antisera against HSP70 showed cytosolic labeling not associated with spheres (**Figure S3D**). The stress granule marker GFP-G3BP1 did not colocalize with cyto-proSAAS (**Figure S3E**). Lysotracker staining also failed to label the structure (data not shown). In addition, the lipid stain Bodipy and antisera to the membrane marker caveolin both failed to co-localize with these structures, indicating that they are not lipid-rich (data not shown). Collectively, these data support the idea that cyto-proSAAS forms membraneless condensates.

### Cyto-proSAAS expression assists the accumulation of TDP-43^216-414^ aggregates

Since cyto-proSAAS efficiently captured GFP-TDP-43^216-414^ aggregates, we examined the effect of cyto-proSAAS on the kinetics of TDP-43 degradation. We used cycloheximide to disrupt protein synthesis during a 6 h period following co-transfection of GFP-TDP-43^216-414^ with cyto-proSAAS; Western blotting was employed to examine the level of cellular GFP-TDP-43^216-414^ forms over time. Interestingly, the half-life of intact GFP-TDP-43^216-414^ was significantly increased upon co-expression of cyto-proSAAS (**Figure S4A**). More than 80% of the GFP-TDP-43^216-414^ protein was degraded after 6 h, whereas 43% of this protein was still present following co-expression with cyto-proSAAS at this time point (**Figure S4A**). Cells not treated with cycloheximide showed similar levels of GFP-TDP-43^216-414^ across the incubation time frame (data not shown). **Figure S4B** shows representative confocal images of HEK cells expressing GFP-TDP-43^216-414^ with or without cyto-proSAAS and/or cycloheximide treatment, showing the cellular persistence of GFP-TDP-43^216-414^ sequestered inside cyto-proSAAS spheres after cyloheximide treatment.

To obtain quantitative and qualitative comparisons of the characteristics of GFP-TDP-43^216-414^ aggregates in the core *vs* the exterior of the proSAAS spheres, we analyzed Z-stacks of these cells (**Figure S4C-E**). We observed that the surface area to volume ratio of GFP-TDP-43^216-414^ aggregates found inside the spheres was significantly larger (i.e. more loosely packed/dispersed) than that of aggregates found outside the spheres (**Figure S4C**). We also noted that the internal GFP-TDP-43^216-414^ fluorescence, both described as mean pixel intensity (**Figure S4D**) and total signal intensity (**Figure S4E**), was significantly lower than that found in aggregates surrounding the spheres. The reduced compactness could indicate a conformational change of sequestered GFP-TDP-43^216-414^ as a result of its interaction with cyto-proSAAS molecules present in the sphere core. We interpret this reduced compactness as possible evidence that GFP-TDP-43^216-414^ aggregates are remodeled within the sphere core.

### Truncated cytoplasmic TDP-43 C-terminal constructs enter cyto-proSAAS sphere cores

To identify the specific region of TDP-43 responsible for its entry into cyto-proSAAS spheres, we overexpressed four additional fluorescent TDP-43 constructs together with cyto-proSAAS, each consisting of different protein domains (see scheme in **Figure 3A**). Similar to TDP-43^216-414^, the C-terminal fragments TDP-43^86-414^ (**Figure 3B**) and TDP-43^170-414^ (**Figure 3C**) both formed cytosolic aggregates which were efficiently recruited into cyto-proSAAS spheres. Interestingly, the C-terminal TDP-43 constructs contain the prion-like domain ^31^. In contrast, the fluorescent N-terminal domain constructs GFP-TDP-43^1-193^ and GFP-TDP-43^1-274^ were concentrated within the nucleus, did not form aggregates, and were not recruited to cyto-proSAAS spheres (data not shown). Since N-terminal-domain-containing TDP-43 constructs were not present in the cytoplasm, we cannot estimate the potential effects of this domain on proSAAS internalization; however, we can conclude that C-terminal domain constructs exhibit strong affinity for entry into cyto-proSAAS spheres.

**Figure 3.**
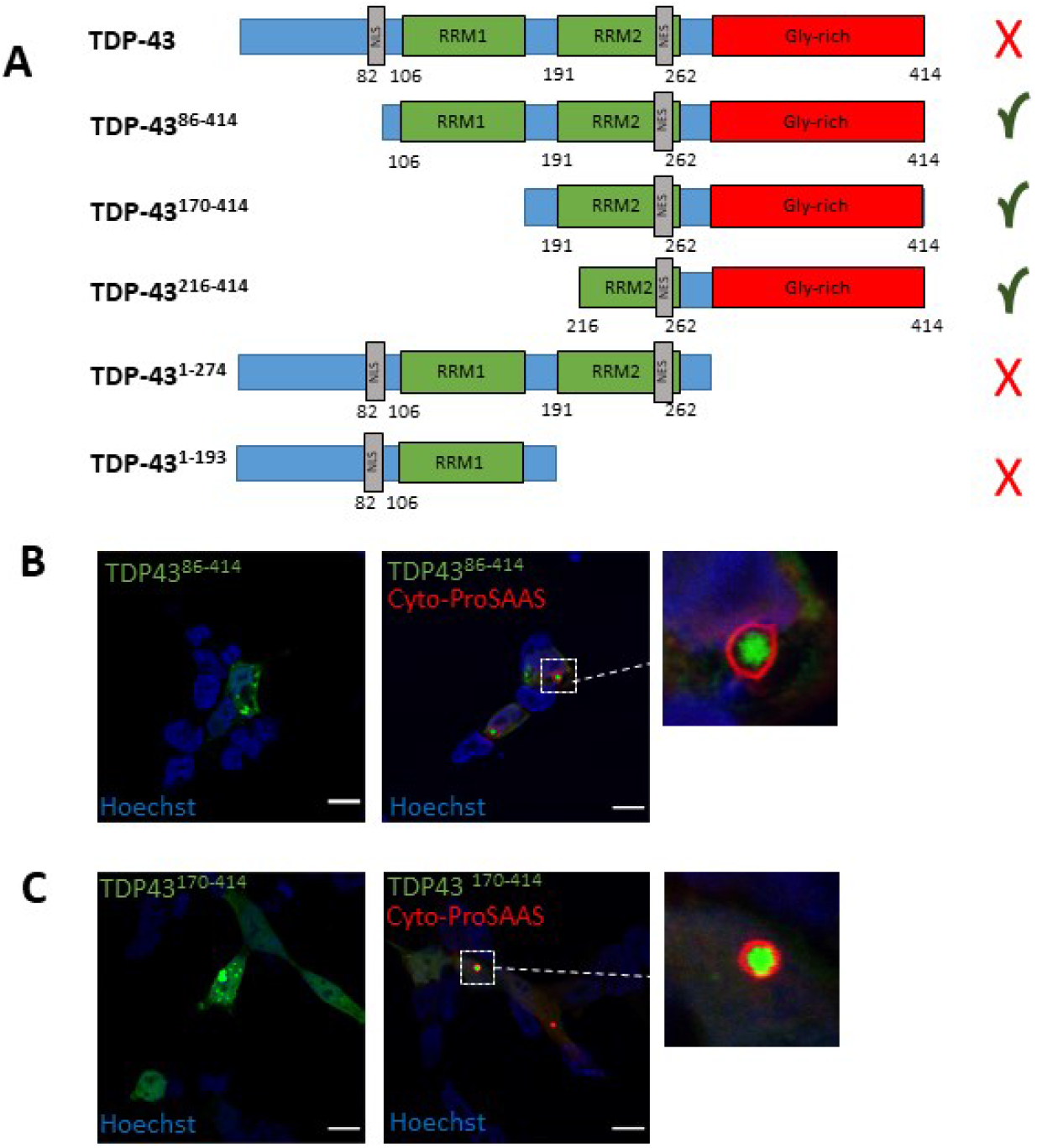
Only TDP-43 fragments lacking the first 85 residues are sequestered by cyto-proSAAS spheres. **Panel A:** Schematic view of TDP-43 constructs. **Panels B and C**: GFP-TDP-43^86-414^ (**panel B**) and GFP-TDP-43^170-414^ (**panel C**) formed aggregates when expressed alone in HEK cells, and these were efficiently incorporated into cyto-proSAAS spheres when this protein was co-expressed. PrLD, prion-like domain; RRM, RNA binding domain; NES, nuclear export signal; NLS, nuclear localization signal. Scale bar, 10 μm. Magnified inserts correspond to 10 μm.

### Conserved amino acids present in the 163-189 segment and a strongly predicted α-helix, residues 37-70, both contribute to efficient cyto-proSAAS sphere formation and to TDP-43^**216-414**^ sequestration

To understand the structural requirements for proSAAS sphere formation we performed structure-function experiments using deletion constructs. ProSAAS is rich in relatively hydrophobic amino acids such as alanine (18.7%) and proline (14.7%) as well as the positively-charged amino acids arginine (13.8%) and lysine (12.4%). ProSAAS also contains several low complexity regions (LCRs) and intrinsically disordered domains (IDDs) throughout the length of the protein, as identified by PLAtform of TOols for LOw Complexity using the SEG tool ^32^ and the Predictor of Natural Disordered Regions tool, respectively (**Figure S5**).

Based on these predictions, we generated a variety of FLAG-tagged cyto-proSAAS constructs lacking specific segments of amino acids within the IDDs and LCRs: 1-30, 37-70, 78-98, 113-134, 163-185, 181-225, and 190-225 (**Figure 4A, S5A**). Constructs that retained the ability to form spheres exhibited either punctate (Δ1-30, Δ113-134, Δ190-225) or diffuse (Δ37-70, Δ113-134) cytoplasmic immunoreactivity, suggesting different morphologies (**Figure 4A, 4C**). Most importantly, we found that the loss of segments 163-185 and 181-225 completely abolished the ability of proSAAS to form spheres (**Figure 4A, 4C)**, whereas the loss of residues 190-225 did not. Taken together, these data support the idea that proSAAS residues 163-189, containing a portion of the conserved region (residues 158-169) ^16^ as well as the highly positively charged site Arg^181^-Arg-Leu-Arg-Arg^185^, strongly contribute to the intermolecular and/or intramolecular interactions that enable sphere assembly.

**Figure 4.**
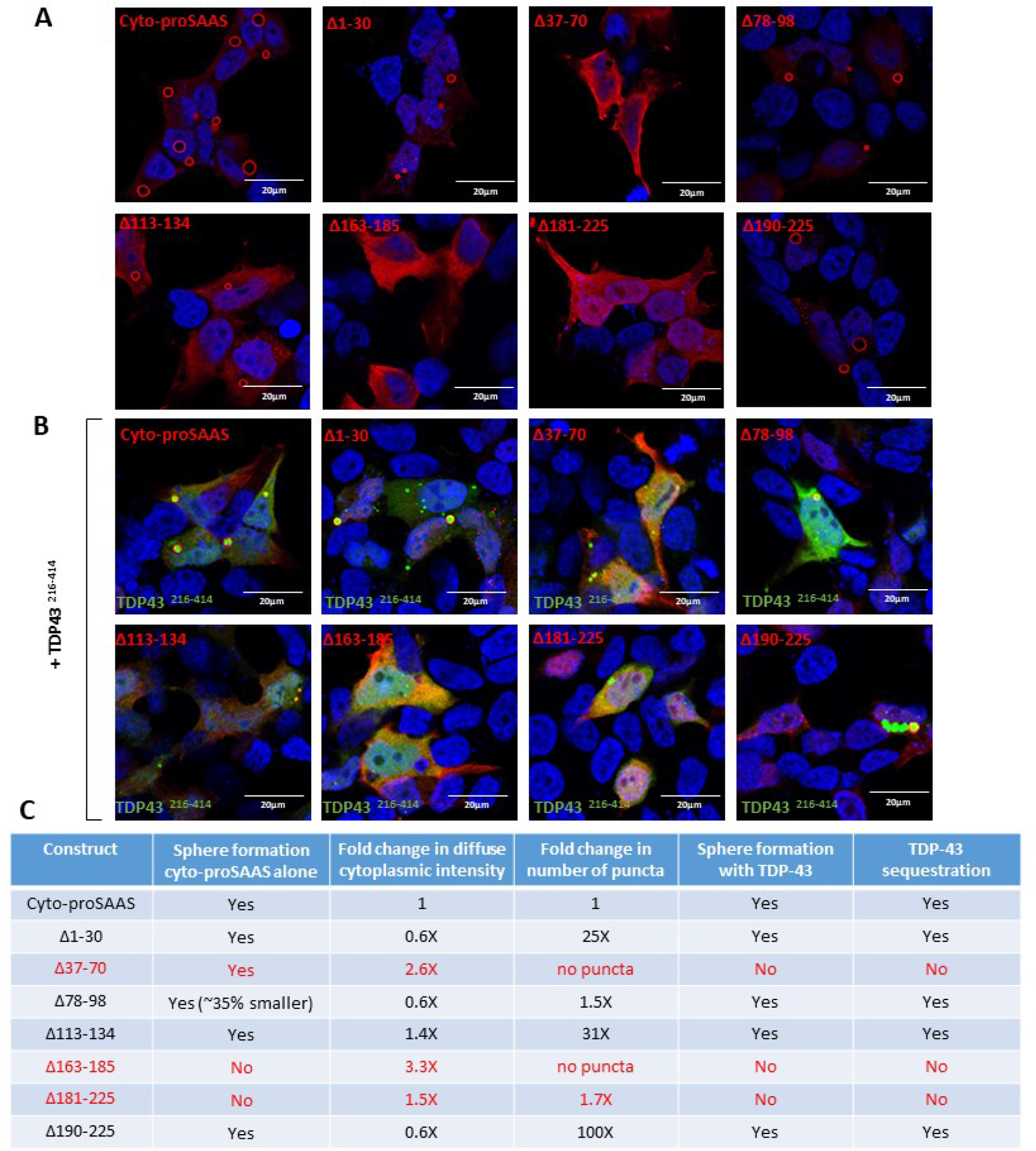
ProSAAS residues 163-189 and 37-70 are required for sphere formation and for GFP-TDP-43^216-414^ sequestration. **Panel A**. Representative images of various deletion constructs of proSAAS expressed in HEK cells showing differences in proSAAS-immunoreactive morphology, ranging from diffuse to punctate to spherical. Scale bar, 20 µm. **Panel B**. Representative images of proSAAS deletion constructs co-expressed with GFP-TDP-43^216-414^ showing differences in their ability to sequester GFP-TDP-43^216-414^. Scale bar, 20 µm. **Panel C**. Summary of results

Our next goal was to assess whether these same deletion constructs could also sequester GFP-TDP-43^216-414^. We found that most constructs that retained the ability to form spheres were also able to successfully encapsulate GFP-TDP-43^216-414^ (**Figure 4B, 4C**). One exception was the cyto-proSAAS Δ37-70 construct. When expressed alone, this construct showed diffuse staining but efficiently formed spheres; but when coexpressed with GFP-TDP-43^216-414^, it failed to form spheres and was unable to encapsulate GFP-TDP-43^216-414^ (**Figure 4B, panel 3; Figure 4C**). GFP-TDP-43^216-414^ coexpression with this construct was also diffuse rather than punctate. According to the recent AlphaFold simulated structure (https://alphafold.ebi.ac.uk/entry/Q9QXV0), residues 37-70 are housed within a strongly predicted coil (32-80) rich in positively (arginine) and negatively (glutamate) charged residues as well as in hydrophobic residues (alanine and leucine). In contrast, the cyto-proSAAS Δ113-134 construct, which showed punctate and diffuse localization, still retained the ability to form spheres and to sequester GFP-TDP-43^216-414^ (**Figure 4B, panel 5; Figure 4C**). In GFP-TDP-43^216-414^ coexpression experiments, constructs lacking the furin site (**Figure 4B:** 163-185, **panel 6**; 180-225, **panel 7**) again did not form spheres, and hence did not sequester GFP-TDP-43^216-414^. In summary, this structure-function analysis shows that the N-terminal 30 residues, the middle 78-98 residues, and the C-terminal 190-225 residues are completely dispensible for sphere formation and GFP-TDP-43^216-414^ sequestration, whereas residues 37-70 and 163-189 are required.

To examine potential effect of proSAAS on TDP-43 aggregation kinetics, we performed *in vitro* assays using recombinant proSAAS. We obtained the TDP-43-His-MBP construct ^33^ which contains a tobacco etch virus (TEV) protease cleavage site between TDP-43 and maltose binding protein (MBP), a protein tag that greatly favors TDP-43 solubility (**Figure 5A**). Upon TEV cleavage, TDP-43 undergoes spontaneous aggregation which can be kinetically measured as absorbance over time ^34^. We found that an excess molar concentration of proSAAS enhanced TDP-43 aggregation kinetics as compared to the control protein bovine serum albumin (BSA) (**Figure 5B**). These data support the cell culture results in showing that proSAAS assists TDP-43 aggregation. However, in preliminary phase contrast experiments we were unsuccessful in observing sphere formation in *in vitro* experiments.

**Figure 5.**
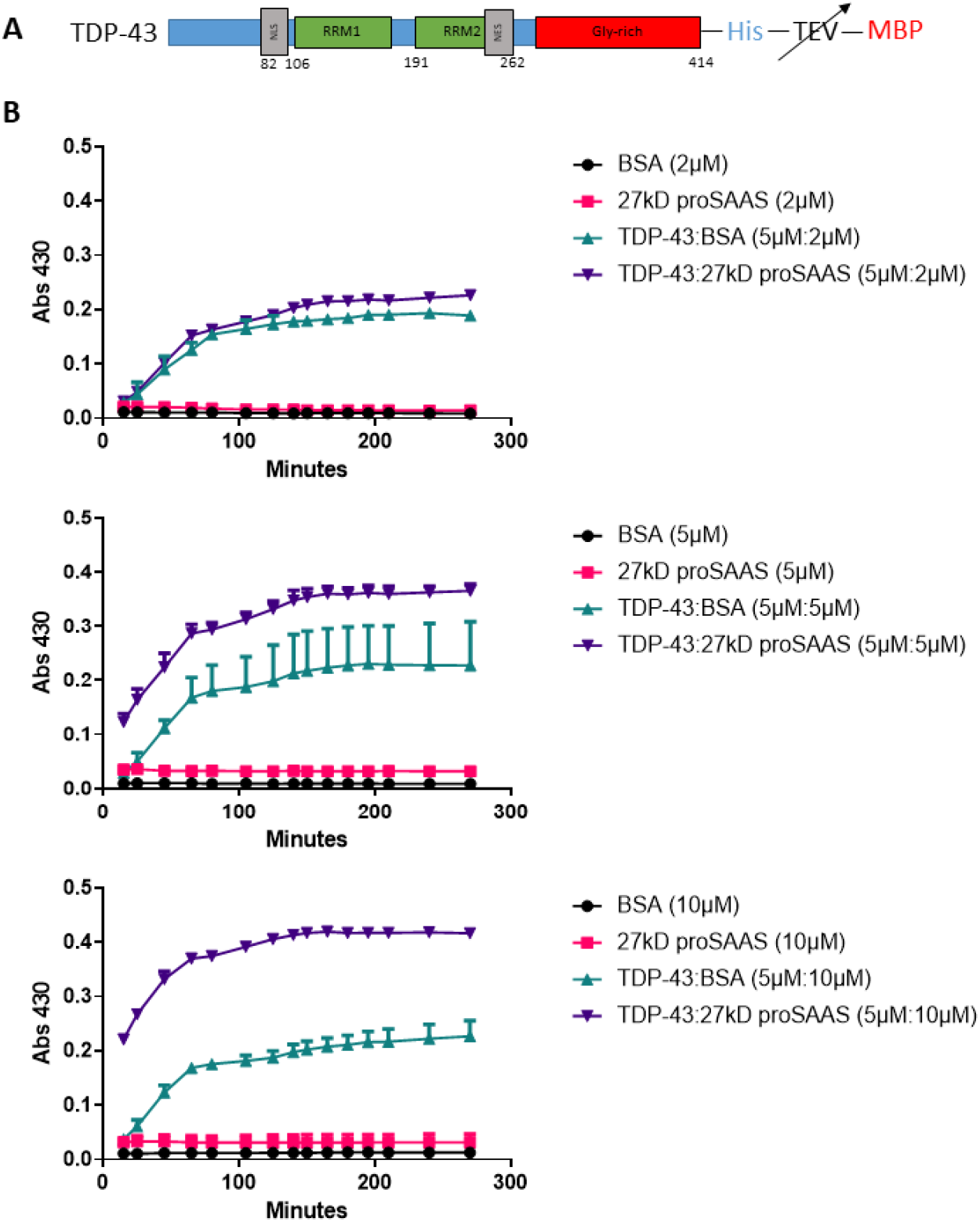
ProSAAS enhances TDP-43 *in vitro* aggregation. **Panel A**: The full-length TDP-43 construct C-terminally tagged with MBP; cleavage of the fusion protein by TEV results in TDP-43 aggregation. **Panel B:** Graphs depicting the absorbance of proSAAS/BSA alone; or mixtures of TDP-43 and proSAAS/BSA at 5 μM: 2 μM (*top*); 5 μM: 5 μM (*middle*); and 5 μM: 10 μM (*bottom*) ratios at 450 nm. ProSAAS addition at a 5 μM: 10 μM ratio of TDP-43 to proSAAS greatly increases TDP-43 turbidity relative to the same molar quantity of BSA.

### Cyto-proSAAS expression is protective against TDP-43 cytotoxicity

Lastly, we examined the cytoprotective effects of proSAAS and another neural and endocrine chaperone, 7B2 ^35^, in a yeast model of TDP-43 toxicity (**Figure 6**). Genetic modifiers identified in yeast have translated well to higher organisms, for example in the identification of ataxin-2 knockdown as a potential therapeutic approach for ALS ^36^, now in clinical trials. In this model, expression of the full-length TDP-43 and proSAAS transgenes is tightly controlled by a galactose-inducible promoter (pGAL1). The empty vector control showed strong TDP-43-mediated growth impairment on galactose agar after 48 h, whereas this toxicity was partially mitigated by expression of proSAAS, cyto-proSAAS, and 7B2. These data provide functional evidence for cytoprotection against full-length TDP-43 toxicity by proSAAS.

**Figure 6.**
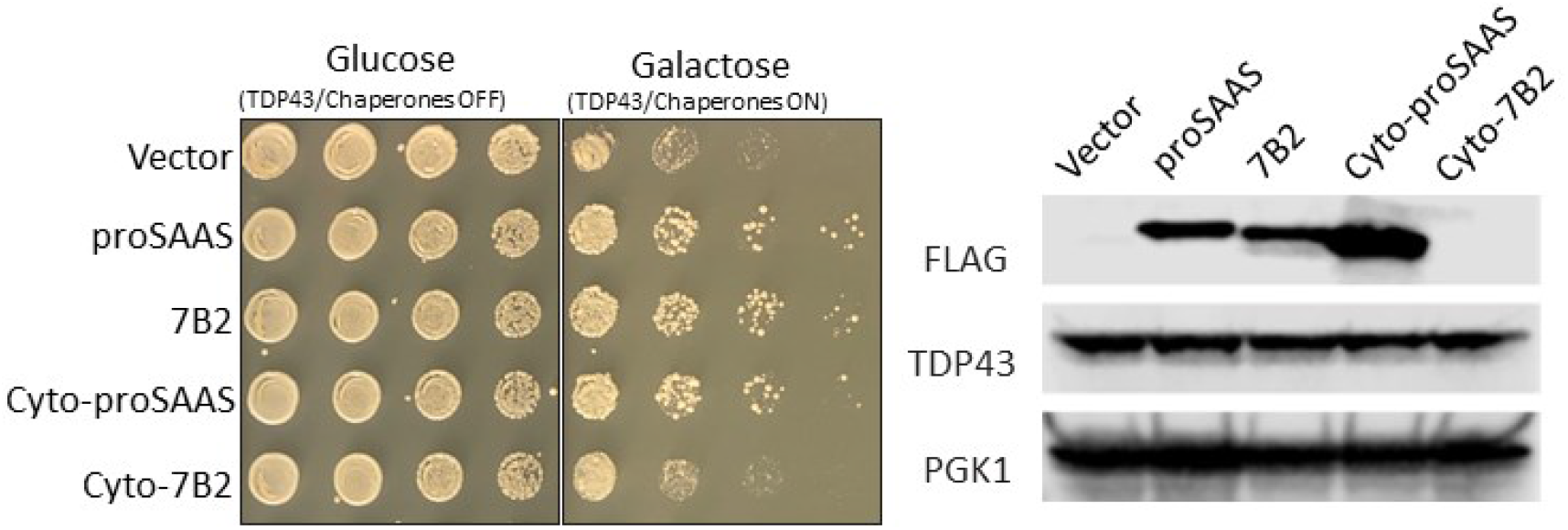
Cyto-proSAAS expression rescues from TDP-43 cytotoxicity in a yeast model. A serial dilution spotting assay of yeast expressing full-length TDP-43 and the indicated chaperone from a galactose-inducible promoter. The empty vector is used as a negative control for chaperone activity and shows TDP-43-induced growth impairment on galactose. **Left panel**: The left side of the spotting plate shows equal spotting and no growth impairment of yeast on glucose media, which does not induce TDP-43 or chaperone expression, while the right side shows growth phenotypes upon induction with galactose. Spotting data are representative images of four independent trials. **Right panel**: The expression of TDP-43 and chaperones was detected via Western blotting; PGK1 is used as a loading control.

## DISCUSSION

Prior work has shown that proSAAS, a small unglycosylated secreted chaperone similar in some aspects to heat shock chaperones ^35^, but which lacks an *α*-crystallin domain, is able to block and/or prevent the aggregation of proteins directly involved in neurodegenerative disorders, including Abeta and *α*-Syn ^15, 16^ ^18 17^. ProSAAS, which is predominantly expressed by neurons and endocrine cells, is known to be associated with plaques, Lewy bodies, and other brain aggregates in AD, PD and FTLD ^15-17^.

In this study, we present data showing cytoplasmically-expressed proSAAS forms spheres that are capable of attracting and sequestering TDP-43 aggregates. These cyto-proSAAS spheres are compact, dense, antibody-impermeable and formed by progressive fusion of smaller spheres, as clearly shown by dynamic imaging. Our finding that cyto-proSAAS spheres do not colocalize with Golgi, lysosomal, membrane and lipid markers supports the idea that the cyto-proSAAS protein spontaneously phase-separates within the cytoplasm. We initially believed that cyto-proSAAS spheres might constitute a form of stress granule; however, arsenite exposure did not increase their formation, nor were we able to identify any surface association of stress granule markers using traditional reporters for the stress-granule associated RNA-binding proteins, G3BP1 or PABPC4. While we confirmed the homogenous presence of cyto-proSAAS within the sphere core using fluorescently-tagged proSAAS, whether cyto-proSAAS spheres in fact also contain other cytoplasmic proteins is presently unclear due to the inability of antibodies to penetrate the sphere shell.

It is now well established that disordered regions, including disordered prion-like domains, play critical roles in phase transitions that lead to the formation of biomolecular condensates (reviewed in ^7, 37^). The unusual phase-separating properties of proSAAS are likely supported by specific intrinsically disordered domains (IDRs) within the protein. While proSAAS does not contain a prion-like domain as predicted by the PLAAC algorithm ^38^, it does contain several predicted IDDs and LCRs (**Figure S5**). Interestingly, we found that both its amino and carboxy-terminal IDRs were dispensible for sphere formation. These deletion studies implicate the predicted coil containing residues 37-70, and suggest the likely involvement of the conserved residues 158-169 and the positively charged sequence 181-185 in sphere formation. Interestingly, several deletion mutants produced small puncta, indicating reduced self-assembly, inefficient fusion and/or complete abrogation of LLPS. Thus, while much of the core of the proSAAS sequence (residues 31-185) appears to contribute to efficient sphere formation, smaller punctal assemblies can be formed from portions of proSAAS.

Surprisingly efficient entrapment of cytoplasmic GFP-TDP-43^216-414^ aggregates occurred within the cores of proSAAS spheres. To our knowledge, no similar cytoplasmic TDP-43-containing particles have been previously reported (reviewed in ^39, 40^). Screening of various fluorescently-tagged potential client proteins indicated that only TDP-43 bears affinity for the cyto-proSAAS core; cyto-proSAAS spheres did not encapsulate fluorescent forms of HTT, tau, *α*-Syn, or other non-aggregating cytosolic proteins, including mCherry. The biochemical basis for the affinity of cyto-proSAAS for TDP-43 is not clear, though likely involves the glycine- and asparagine-rich prion-like domain within the carboxy-terminal domain, thought to represent a major determinant for aggregation ^40-42^. The structure-function studies presented here show that TDP-43 constructs containing the 86-residue N-terminal domain are not recruited to cyto-proSAAS spheres, whereas all longer TDP-43 constructs are encapsulated. We suspect that this phenomenon is due to the lack of abundance of these nuclear-localized proteins within the cytoplasm rather than to direct exclusion. Dynamic imaging of cyto-proSAAS spheres containing fluorescent TDP-43^216-414^ aggregates revealed that cyto-proSAAS spheres containing GFP-TDP-43^216-414^ cargo undergo continuous fusion and enlargement; and that fluorescence derived from internalized GFP-TDP-43 aggregates is more dispersed (as measured by fluorescence intensity divided by area) than is external GFP-TDP-43 fluorescence, suggesting interaction of internal cyto-proSAAS with GFP-TDP-43^216-414^.

Physiological cellular spherical condensates of 2-4 μm have to our knowledge not yet been reported. Most cellular condensates reported to date are nuclear and small, less than one micron in diameter (reviewed in ^7 37 39^). Recent studies have employed synthetic condensates to understand the role of phase separation in cellular processes ^43^. Remarkably, the phase-separating bacterial protein PopZ, which has been used to construct short (76-residue) engineered “PopTags” for targeting specific cellular proteins to condensates ^44, 45^ contains a sequence, DVVRELLRPLLKEWL, which is highly homologous to the DVDPELLRYLLGRIL sequence (residues 153-167) within proSAAS (60% identical). Interestingly, this specific segment represents one of the most conserved regions within proSAAS ^16, 46^, and preliminary experiments indicate that it is essential to sphere formation (results not shown). This finding leads us to speculate that a proSAAS-related sequence might conceivably also be employed to construct a mammalian PopTag; in this scenario, cyto-proSAAS could be engineered as a designer “holdase” condensate to reduce the concentration of cytoplasmic aggregates (see also ^47^).

Indeed, synthetic condensates involving TDP-43 aggregates have also recently been reported. Yu *et al* have found that an acetylated, non-RNA-containing form of TDP-43 is able to form dense spherical nuclear annuli (“iLSA”) which entrap Hsp70-type chaperones ^48^. These TDP43/Hsp70 spheres differ from cyto-proSAAS spheres in several important respects: they require the presence of an energy-dependent chaperone to drive sphere formation; they require the N-terminal domain of TDP-43; and they contain structured assemblies of TDP-43 species as an outer shell for internalized Hsp70, while apparently unstructured TDP-43^216-414^ aggregates are sequestered within the cores of cyto-proSAAS spheres. Aromatic residues play a key role in TDP-32 LLPS ^49^. The identification of specific residues within TDP-43^216-414^ that underlie its affinity for chaperones, including proSAAS spheres, will representan important step towards manipulating the aggregative properties of this clinically important protein.

Cyto-proSAAS sphere expression clearly results in a variety of functional consequences. Cyto-proSAAS expression increases the half-life of TDP-43^216-414^, most likely reflecting the sequestration of TDP-43^219-414^ aggregates. We were unable to assess potential protective effects of cyto-proSAAS on TDP-43-mediated cytotoxicity in mammalian cells simply because TDP-43 expression was not reliably cytotoxic in any system we tested, including PC12 cells. However, in yeast cells, cyto-proSAAS partially rescued the growth impairment caused by expression of full-length TDP-43. Previous work on potentiated Hsp104 disaggregases show that Hsp104-mediated rescue of aggregate toxicity in yeast models can be replicated in an animal nervous system ^50^. Other modifiers of TDP-43 toxicity in yeast have also translated well to mouse models ^36, 51^. Thus, our findings in yeast are highly promising in their suggestion of cytoprotective effects for proSAAS in higher eukaryotes.

In summary, we have here identified a novel mechanism for aggregate protein sequestration via proSAAS sphere encapsulation. We suggest that further studies exploring the active elements within cyto-proSAAS which interact with TDP-43 will be useful in designing pharmacologic agents - and possibly synthetic condensates – which are able to enhance aggregate sequestration.

## EXPERIMENTAL PROCEDURES

### Cell culture

Neuro2A and HEK cells were obtained from ATCC (Manassas, VA) and grown in OptiMEM/DMEM and DMEM, respectively, both with 10% fetal bovine serum (FBS; Gemini Bio, West Sacramento, CA). Transfection of cells was carried out on 70-80% confluent cells in a 24-well plate fitted with poly-L-lysine-treated glass coverslips, using 0.5 - 1 μg of cDNA and FuGene as per the manufacturer’s instructions (Promega, Madison, WI). Cells were fixed for immunocytochemistry 24 or 48 h after transfection, as stated.

Hippocampal primary neurons obtained from E17 rat embryos were cultured as described previously ^52^ using digestion with 0.25% trypsin-EDTA and mechanical dissociation by gentle pipetting through a series of small-bore Pasteur pipettes. Filtered cells were resuspended in Neurobasal medium (InVitrogen) supplemented with 5% B27 supplement, 5% penicillin-streptomycin, and 2 mM L-GlutaMAX (all from Invitrogen), and then cultured on poly-L-lysine coated 12-mm coverslips at 37 °C and 5% CO2 in a humidified atmosphere. Media were replaced every 3-4 days. Seven-day old primary hippocampal cells, grown on poly-L-lysine coated 12 mm coverslips, were transfected with 2 μg of cyto-proSAAS cDNA using Lipofectamine (Invitrogen). Cells were fixed, immunostained with FLAG and HA tag antisera, and imaged 24 h post-transfection. Between 5-10% of cells were transfected by this method.

### Expression vectors

The majority of the constructs used for expressing aggregating proteins were obtained from Addgene (Watertown, MA). The Addgene catalog numbers are as follows: GFP-tagged TDP-43 constructs TDP-43^216-414^, #28197; TDP-43^1-193^ (#28202); TDP-43^1-273^ (#28200); TDP-43 ^86-414^ (#28195) and TDP-43^170-414^ (#28196), described in ^53^; HTT-Q74 (HTT exon 1 Q74, His- and HA-tagged, #40264); and GFP-HTT-Q74, #40262, in ^54^; EGFP-Tau, #46904, and EGFP-TauE14, #46907 ^55^; EGFP-α-Syn, #40822 and EGFP-α-Syn-A53T, #40823 ^56^. The stress granule marker GFP-G3BP1 was also obtained from Addgene (#135997 ^57^), while the LC3-mCherry construct was obtained from Dr. Marta Lipinksi (U. Maryland-Baltimore; ^58^).

Murine proSAAS, which is 81% identical to human proSAAS, was used in these studies for consistency with previous work ^16^. A FLAG-tagged eukaryotic expression vector for murine proSAAS was constructed by Genscript (Piscataway, NJ) in pcDNA3.1(-) (hygromycin-resistant; Invitrogen) by insertion of the nucleotide sequence GATTACAAGGATGACGACGATAAG, encoding the DYKDDDDK FLAG peptide, following the signal peptide. Signal-less cyto-proSAAS was created with a forward primer introducing a methionine amino terminal to the FLAG tag, and a reverse primer targeted to the carboxy-terminus of proSAAS. This product was cloned into the *NheI* and *HindIII* sites of the pcDNA3.1(-) hygromycin resistance-encoding vector (Invitrogen). Lastly, the mCherry proSAAS fusion construct was made by N-terminal insertion of mCherry into the signal-less proSAAS construct. A cytoplasmic mCherry expression vector was obtained from Dr. Megan Rizzo (University of Maryland-Baltimore, MD). All construct sequences were confirmed by sequencing.

### Confocal microscopy

For immunohistochemistry (IHC), cells were transfected in a 24-well plate on 12-mm coverslips and fixed in 4% PFA in PBS for 20 min at 24 or 48 h after transfection, as indicated. Cells were incubated for 30 min at room temperature in IHC blocking buffer (PBS containing 5% FBS, 0.5% Triton X-100 and 100 mM glycine). Antibodies against FLAG (Sigma-Aldrich, #F7425), Rab7a (# NBP1-87174, Novus); HSP70 (#11660-T52; Sino Biological), TDP-43^216-414^ (#NB11O-5537; Novus), and GFP (#GFP879484; Aves Labs, Inc) were used at a dilution of 1:1000 in IHC blocking buffer and incubated for 30 min at room temperature. Anti-HA tag antiserum (Abcam, #130275) was incubated with cells overnight at a dilution of 1:200. After washing multiple times with PBS, cells were incubated with secondary fluorescent antibodies (Invitrogen) at a dilution of 1:1000 and Hoechst stain at a dilution of 1:5000. In some experiments, primary antibodies conjugated with fluorescent dyes were used; these were anti-FLAG (Alexa647; #637315; BioLegend), and anti-giantin (Alexa488; #908701; BioLegend), anti-LAMP-1 (Alexa488; #121607; BioLegend). The specificity of the proSAAS signal obtained with anti-FLAG antiserum was confirmed using IgG-purified antibody raised against the His-tagged, recombinant murine 21 kDa proSAAS protein (proSAAS^1-180^) ^15^. Morphological characterization of proSAAS spheres and quantitation of fluorescent signal and transfection efficiency was carried out using Fiji/ImageJ ^59^. At least two 40x fields, obtained from three independent experiments using the same experimental conditions, were combined for statistical purposes.

To study the nature of TDP-43^216-414^ aggregates, Z-stacks (0.2 μm intervals) of individual cells expressing TDP-43^216-414^ aggregates, both within and outside cyto-proSAAS spheres, were obtained. For quantitative imaging, 12-bit images were taken, with care to avoid saturation of the GFP signal. The GFP signal within and outside the proSAAS spheres was quantified using 3D ImageJ Suite ^60^. Images were first smoothed with a 1.5 pixel Gaussian blur and then segmented in the cyto-proSAAS channel with manual threshold adjustment to select only spheres. The TDP-43^216-414^ channel was segmented with a threshold to include only aggregated GFP signal. Binary masks were generated using the segmented images and used to quantify the original GFP signal inside and outside the cyto-proSAAS sphere, using the Measure 3D and Quant 3D plugins.

### STED microscopy

HEK cells were plated on 12 mm coverslips at 70-80% confluency and transfected with sec proSAAS or cyto-proSAAS cDNAs. After 24h, the coverslips were fixed and immunostained using homemade anti-proSAAS antibody as described before and STAR-RED secondary antibody compatible with Stimulated Emission Depletion (STED) microscopy. The Nikon A1RHD25/STED microscope was used to generate super-resolution images of proSAAS spheres.

### Dynamic in vivo imaging

Live cell confocal microscopy was performed on a Nikon W1 spinning disk confocal system, on a Nikon Ti2 inverted microscope, with a Hamamatsu sCMOS camera, at the Confocal Microscopy Core Facility, University of Maryland Baltimore. Acquisition was controlled with Nikon Elements software. Cells were incubated at 37°C and under 5% CO2 in Opti-MEM media during imaging. Cells were grown in high grid 500 35-mm plates (Ibidi; Fitchburg, WI) to 50% confluence, and double-(TDP-43^216-414^ and cyto-proSAAS; equal ratio) or triple-transfected (TDP-43^216-414^, cyto-proSAAS and Cherry cyto-proSAAS; 1, 1, and 0.2 µg respectively) for 6 to 24 h prior to imaging GFP and mCherry fluorescence, as indicated in the figures. Live cell DIC and epi-fluorescent microscopy was performed on an Olympus VivaView. Cells were plated in 35-mm glass-bottomed dishes (MatTek; Ashland, MA) and transfected 18 h prior to the start of imaging. Multiple regions of cells were selected and imaged at 2 min intervals over a period of 6 h, using excitation/emission filter sets for GFP and RFP. Image analysis was performed with ImageJ.

### Electron microscopy

HEK 293 cells were grown on 12-mm coverslips up to 70% confluency and transfected with empty vector, TDP-43^216-414^ cDNA alone, or TDP-43^216-414^ in combination with cyto-proSAAS cDNA. Twenty-four h after transfection, cells were fixed in a solution of 2% paraformaldehyde, 2.5% glutaraldehyde, in 0.1 M PIPES buffer (pH 7). After washing, cells were quenched with 50 mM glycine in 0.1 M PIPES buffer (pH 7) for 15 min and washed again with 0.1 M PIPES buffer, post-fixed in 1% (w/v) osmium tetroxide, 0.75% ferrocyanide in 0.1 M PIPES buffer for 60 min, washed with 0.1 M PIPES, and stained with 1% (w/v) uranyl acetate in water for 30 min. After washing, specimens were dehydrated using serially graded ethanol solutions (30%, 50%, 70%, 90% and 100%) and infiltrated and embedded in Spurs resin (Electron Microscopy Sciences, Hatfield, PA) following the manufacturer’s recommendations. Ultrathin sections (70 nm) were cut on a Leica UC6 ultramicrotome (Leica Microsystems, Inc., Bannockburn, IL), collected onto copper grids, and examined in a Tecnai T12 transmission electron microscope (Thermo Fisher (Formerly FEI. Co.), Hillsboro, OR) operated at 80 kEV. Digital images were acquired by using a bottom-mounted CCD camera (Advanced Microscopy Techniques, Corp, Woburn, MA) and AMT600 software.

### Western blotting

Western blotting of HEK cell extracts was performed using 70-80% confluent cells grown in 12-well plates. Cells were resuspended in 150 μl of RIPA buffer (25 mM Tris pH 7–8, 150 mM NaCl, 0.1% SDS, 1% Triton X-100 and 0.5% sodium deoxycholate, supplemented with protease inhibitors [“Complete”; Sigma-Aldrich, St. Louis, MO]) and frozen prior to assay. Samples were thawed, boiled, electrophoresed on 15% acrylamide gels, and blotted to nitrocellulose using a BioRad Turbo Blotter at 250 V for 10 min. Blots were blocked in 5% blotting grade milk (Bio-Rad) in 0.05% Tween-20 in Tris-buffered saline, pH 7.4 (blotting buffer) and were then incubated with primary antisera against GFP (1:1500; #879484; Aves Labs, Inc), anti-FLAG (Sigma-Aldrich, #F7425), or anti-actin (#A2228; Sigma) in blotting buffer overnight at 4C. Secondary antisera, used at dilutions of 1:5000 in blocking buffer, were HRP-linked (Bio-Rad # s170-6516, 170-6615, and L005680A, respectively).

### Cycloheximide experiments

HEK cells were plated in 12-well plates and transfected when 70-80% confluence was reached. Cycloheximide (20 μg/ml; Sigma-Aldrich) was used to inhibit total protein synthesis 24 h after transfection. The cells were monitored for 6 h following addition of drug, a duration that resulted in 80% reduction of aggregates in cells transfected with GFP-TDP-43^216-414^. Cells were lysed in Laemmli sample buffer at the times indicated in the Results and subjected to blotting using chicken antiserum against GFP (Aves Labs). Three independent experiments were used for quantification purposes. To investigate the cellular distribution of proteins following 6 h of cycloheximide treatment, cells were fixed as previously described, immunostained using anti-FLAG antibody (Sigma-Aldrich, #F7425), and imaged by confocal microscopy.

### Bioinformatics and site-directed mutagenesis for structure-function studies

The cyto-proSAAS amino acid sequence was analyzed using the online tools Predictor of Natural Disordered Regions (PONDR) for intrinsically disordered domains (IDDs) and PLAtform of TOols for LOw Complexity using the SEG tool for low complexity regions (LCRs). Based on this analysis, a series of FLAG-tagged cyto-proSAAS constructs lacking the sequences encoding the amino acids 1-30, 37-70, 78-98, 113-134, 163-185, 181-225 or 190-225 were generated using site-directed mutagenesis and cloned into pcDNA3.1(-) by Genscript (Piscataway, NJ) for eukaryotic expression. All constructs were verified by bidirectional cDNA sequencing. Vectors were transfected either alone or together with a vector encoding GFP-TDP-43^216-414^ into 70-80% confluent HEK cells. Cells were fixed and immunostained with anti-FLAG antiserum (Alexa647; #637315; BioLegend) and Hoechst dye to assess cyto-proSAAS sphere formation and TDP-43 sequestration 24 h after transfection. Mean intensity and sphere dimensions were measured using ImageJ software.

### Protein purification and in vitro TDP-43 aggregation assay

The pJ4M/TDP-43 vector encoding 6xHis-TDP-43-TEV-MBP was obtained from Addgene (#104480; ^33^). A His-tagged proSAAS cDNA (lacking the signal peptide) was cloned into pET45a(-) using a GoTaq® Flexi DNA Polymerase kit (M829). Starter cultures of *E. coli* BL21 DE3 cells, transformed with each construct, were used to inoculate 1 L of autoinduction medium containing NPS (0.025M (NH_4_)_2_SO_4_, 0.05M KH_2_PO_4_, 0.05 M Na_2_HPO_4_) and 50/5/2 (0.5% glycerol, 0.05% glucose and 0.02% lactose) and incubated overnight at 30°C at 210 RPM for expression by autoinduction. The TDP-43 fusion protein was purified from cell extracts using Ni-NTA and amylose affinity chromatography following published protocols ^33, 34^. Recombinant proSAAS was purified by Ni-NTA affinity chromatography, as previously described ^16^.

To perform the *in vitro* TDP-43 aggregation assay, the TDP-43-TEV-MBP fusion protein was first exchanged into aggregation buffer (20 mM Hepes, pH 7.0; 150 mM NaCl, and 1 mM DTT) and centrifuged, as previously described ^34^. Final concentrations of 5 µM TDP-43-TEV-MBP and the indicated final concentrations of recombinant proSAAS or BSA (NEB BioLabs; B9001S) were achieved by mixing aliquots of a 44 μM stock solution of TDP-43-TEV-MBP with aliquots of stock solutions of either 55 μM proSAAS or 151 μM BSA (both in 5 mM acetic acid). These solutions were added to 100 units of TEV protease (NEB) in aggregation buffer, present in black-walled 96-well clear-bottom plates (Costar #3904). The assay was carried out in triplicate in a total volume of 100 μl aggregation buffer; control wells contained only proSAAS or BSA and TEV in the same volume of aggregation buffer. The absorbance was measured at 450 nm using a Biorad plate spectrophotometer every 15 min over 5 h.

### Yeast cell screen for proSAAS/cyto-proSAAS rescue from TDP-43 cytotoxicity

Yeast experiments were performed using the BY4741*Δhsp104* strain. Expression of TDP-43 was induced from the pGAL1 promoter, which is maintained on a HIS3-selectable CEN plasmid (pEB413GAL_TDP-43). ProSAAS and 7B2 (and their signal-less forms) were expressed from pGAL1 on a separate URA3-selectable CEN plasmid (pEB416GAL). Prior to the yeast spotting assays and Western blotting, the yeast were cultured for 12 h at 30°C in pre-induction media containing 2% raffinose + synthetic-defined media lacking histidine and uracil (-his/ura). For spotting assays, the samples were normalized to an OD600 of 2 and serially diluted five-fold per spot. Rescue of TDP-43 cytotoxicity was assessed after 48h of spot growth on 2% glucose or galactose -his/ura agar plates at 30°C. For Western blotting, the yeast were inoculated in liquid induction media containing 2% galactose -his/ura and grown for 6 h to induce protein expression. Normalized cell pellets were treated for 5 min with 0.1 M NaOH followed by resuspension in 1x SDS-PAGE sample buffer (100mM Tris-HCl pH 6.8, 4 mM EDTA, 4% SDS, 0.1% Bromophenol blue, 20% glycerol) + fungal protease inhibitor cocktail (Sigma, P8215) and then heated at 95°C for 10 min. Samples were electrophoresed on a 4-20% Criterion Precast Tris-HCl acrylamide gel (200V for 1 h) and transferred to a PVDF membrane (0.15 amp for 1 h). Blots were blocked with Li-COR Odyssey Blocking Buffer and then incubated with primary antibodies: anti-FLAG (Sigma F1804, 1:1000 for proSAAS and 7B2), anti-TDP-43 (Proteintech 10782-2-AP, 1:1000), or anti-PGK1 (Invitrogen 459250, 1:1000). Blots were visualized after a 30 min incubation with secondary antibodies (Li-COR 680RD anti-rabbit 1:2500; Li-COR 800CW anti-mouse 1:5000).

### Statistical analysis

We used GraphPad Prism 5.0 and 6.0 (GraphPad Software, La Jolla, USA) for statistical analyses and data presentation. Graphs represent means ± SD or SE, as indicated, with the number of replicates indicated in the Methods or Figure Legends. Student’s unpaired two-tailed t-test was used for comparison of data obtained for fluorescence density measurements from at least three independent experiments, with p<0.05 considered significant.

## Acknowledgments

We are grateful for research support from NIH grant AG 062222 to IL and support from “*Convocatorias de movilidad”* from University of Castilla-la Mancha (UCLM) to JRP. This work utilized an EM sample preparation instrument that was purchased with funding from a National Institutes of Health SIG grant (1S10RR26870-1). We are grateful for the assistance of Dr. Ru-ching Hsia of the Electron Microscopy Core Imaging Facility, Center for Innovative Biomedical Resources, University of Maryland-Baltimore. We acknowledge the use of the Confocal Microscopy Core Facility, University of Maryland-Baltimore. We thank Ms. Minerva Contreras for hippocampal cells, and our UMB colleagues Drs. Rizzo and Lipinski for various constructs. We thank Dr. Bede Portz for help with the MBP-TDP43 aggregation assay. EB was supported by a Milton Safenowitz Post-Doctoral Fellowship from ALSA and by NIH grant F32NS108598. JS was supported by grants from ALSA, Target ALS, and The Robert Packard Center for ALS Research at Johns Hopkins.

## Requests for resources and reagents

Requests should be directed to Iris Lindberg.

