## Supplementary figures for "Sequestration of TDP-43^216-414^ aggregates by cytoplasmic expression of the proSAAS chaperone"

**Supporting Information (5 figures)**

**Supplemental Figure S1. Cyto-proSAAS forms spheres in various cell types, serving as a potential platform to identify encapsulated cytoplasmic targets.**

**Panel A:** Representative confocal images of Neuro2A cells transfected with cyto-proSAAS cDNA for either 24 h (*left*) or 48 h (*right*) showing characteristic Flag-immunoreactive spheres (shown enlarged at right). **Panel B:** Overexpressed cyto-proSAAS forms spheres that can be observed using phase-contrast microscopy (*left side*, **yellow arrows**) and which consists of mostly unprocessed 30 kDa proSAAS, observed as a band in a Western blot using FLAG antibody (*right side*). **Panel C:** Expression of cyto-proSAAS in primary rat hippocampal cells results in the formation of spheres similar to those observed in Neuro2A and HEK cells. The border of the transfected cell is outlined in white. **Panel D:** Expression of Cherry-cyto-proSAAS (*red*) generates dispersed cytosolic aggregates in Neuro2A cells. **Panel E:** Representative images of GFP-tagged Tau; Tau E14; $\alpha$-synuclein; $\alpha$-synuclein A53T; TDP-$43^{216-414}$; HTT-Q74; and HA-tagged HTT-Q74. Images were taken at 40X at 24h after transfection. Scale bar, 10 mm. Magnified inserts correspond to 10 mm.

Figure S1

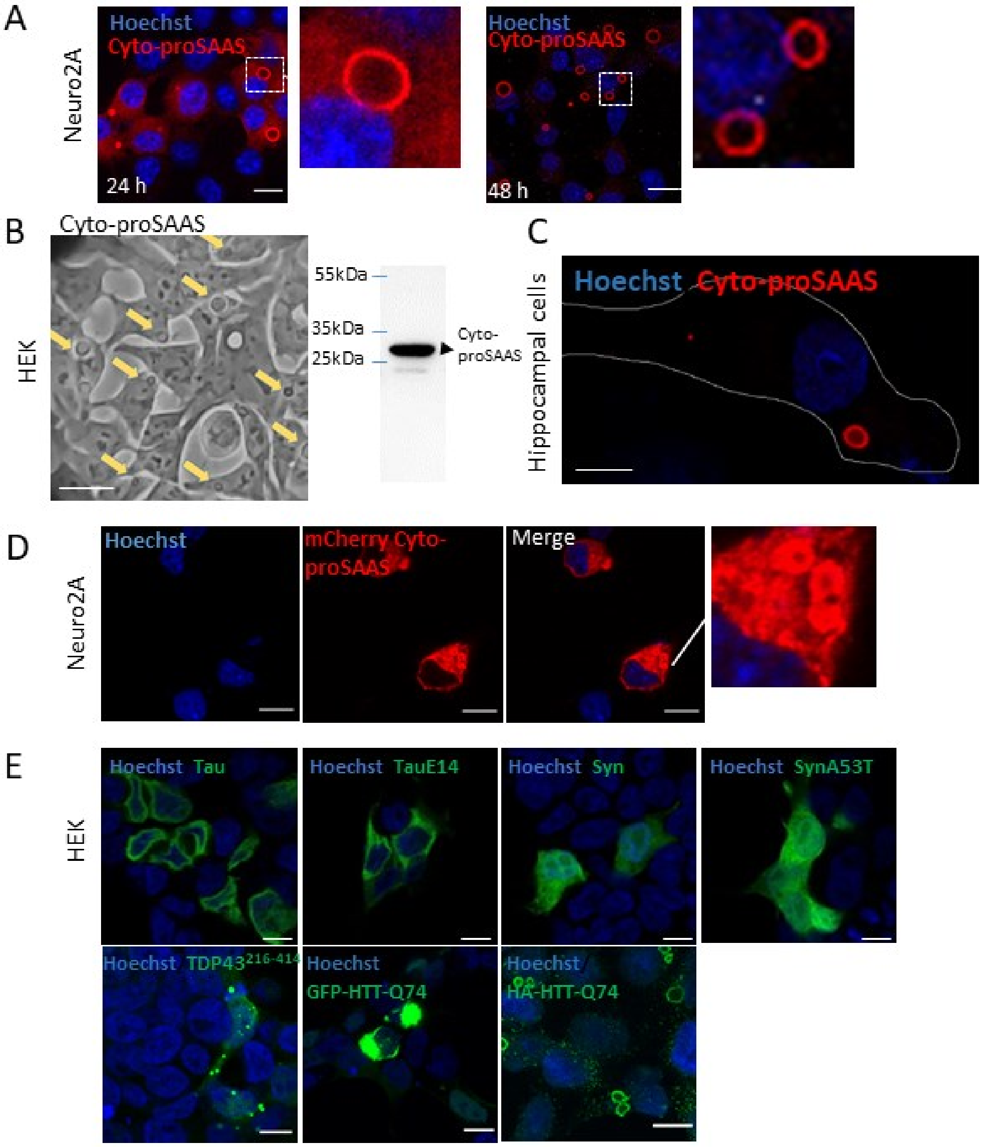

**Supplemental Figure S2. Cyto-proSAAS spheres specifically encapsulate TDP-43$^{216\text{-}414}$ aggregates, but not HTT-Q74.**

**Panel A** Cyto-proSAAS expression alone. **Panel B:** TDP-43$^{216\text{-}414}$ aggregates**,** but not HTT-Q74 aggregates (**panel C**) nor GFP-HTT-Q74 aggregates **(panel D)** are found inside cyto-proSAAS spheres (*red*) in HEK cells. **Panel E** shows the characteristic cytosolic fluorescence of dispersed mCherry (*red*) aggregates that never become accumulated within cyto-proSAAS spheres (*green*). Images were taken at 40X 24 h after transfection. Scale bar, 10 μm. Magnified images correspond to 10 μm.

Figure S2

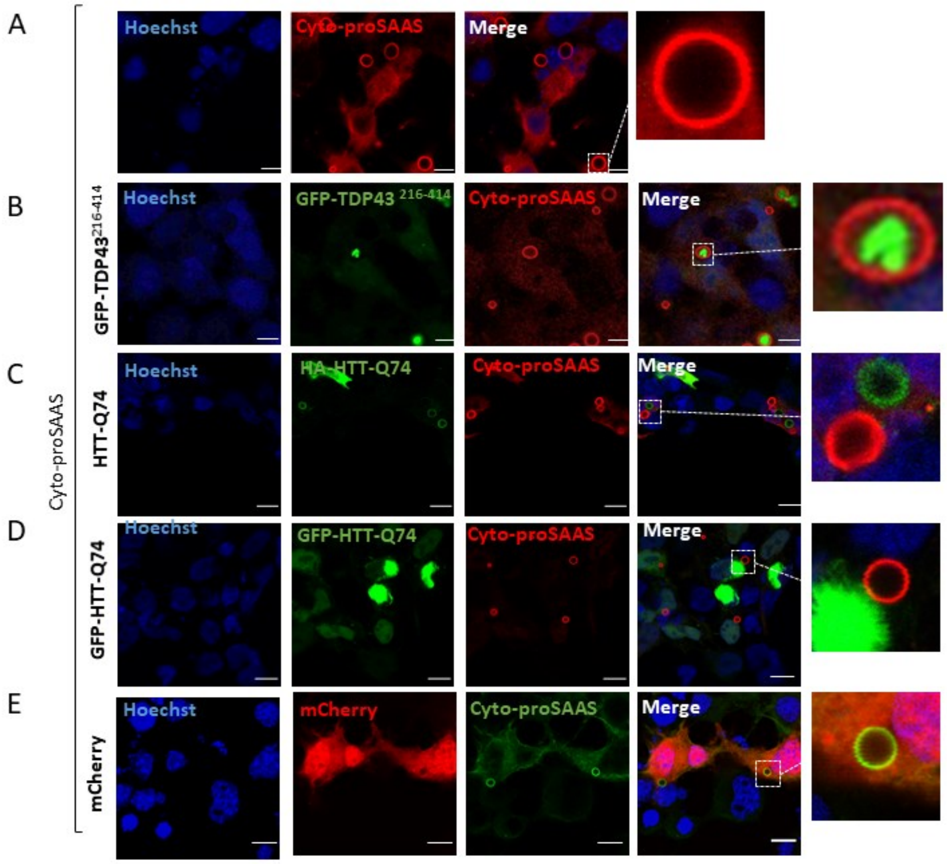

**Supplementary Movie 1. Dynamic imaging of mCherry-cyto-proSAAS spheres showing fusion of large spheres.** Dynamic widefield microscopy of mCherry-cyto-proSAAS (*red*), beginning 18 h post transfection of cyto-proSAAS and mCherry-cyto-proSAAS (1: 0.1 ratio).

**Supplemental Movie 2. Dynamic imaging of mCherry-cyto-proSAAS spheres showing sphere fusion with simultaneous GFP-TDP-43$^{216\text{-}414}$ sequestration.**
Movie of **Figure 2D**. Dynamic widefield microscopy of mCherry-cyto-proSAAS *(red*) and GFP-TDP-43$^{216\text{-}414}$ (*green*), and DIC (*grayscale*) beginning 18 h post transfection of cyto-proSAAS, GFP-TDP-43$^{216\text{-}414}$ and mCherry-cyto-proSAAS (1:1:0.2 ratio). Each frame corresponds to 2 min in real time. The ImageJ plugin Linear Stack Alignment with SIFT was used to stabilize the movie around the transfected cell.

**Supplemental Figure S3. Cyto-proSAAS spheres are not associated with cellular markers of autophagy, lysosomes, Golgi, cell stress, and stress granules.**

**Panel A:** No fluorescent signal was observed associated with cyto-proSAAS spheres following triple transfection of LC3-Cherry, cyto-proSAAS and GFP-TDP-43$^{216\text{-}414}$ cDNAs.

**Panels B-D**: Lack of cyto-proSAAS sphere immunostaining with antisera against Lamp-1 (**panel B**) giantin (**panel C**) and HSP70 (**panel D**).

**Panel E:** Cyto-proSAAS neither associates with nor sequesters the co-expressed stress granule marker GFP-G3BP1.

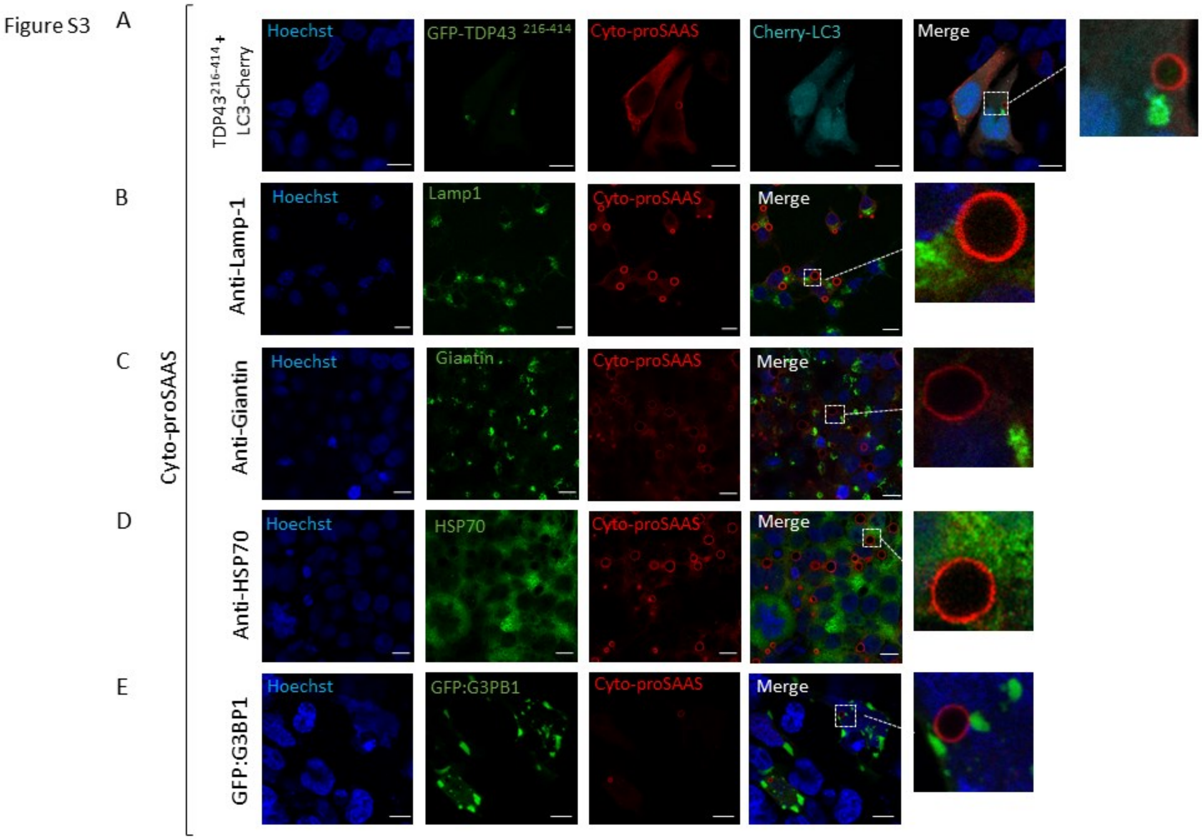

**Supplemental Figure S4. Co-expression of cyto-proSAAS increases the cellular half-life of GFP-TDP-43$^{216-414}$.**
**Panel A:** HEK cells transfected with cDNAs encoding GFP-TDP-43$^{216-414}$ alone, or in combination with cyto-proSAAS, were monitored by Western blotting using an anti-GFP antibody (*left panels*). Quantitation of triplicates is shown in below the blot. Approximately 80% of GFP-TDP-43$^{216-414}$ disappears after 6 h of cycloheximide treatment when cells are co-transfected with an empty vector; however, 43% remains in cells co-transfected with cyto-proSAAS cDNA (*, $p<0.05$).
**Panel B:** Confocal images taken prior to and following cycloheximide treatment confirm reduced cellular quantities of GFP-TDP-43$^{216-414}$ following cycloheximide treatment in the absence of cyto-proSAAS expression (*left panels*); in contrast, TDP-43$^{216-414}$ fluorescence is retained within cyto-proSAAS spheres (*right panels*).
**Panels C-E:** Ratio of the total surface area of GFP-TDP-43$^{214-416}$ to the total volume of the same objects (**C**); the mean pixel intensity for each group (**D**);the total signal intensity of each group (**E**), determined by the sum of signal intensity of all pixels in that group. (n=12 cells, two-tailed paired t-test *$p<0.05$, **$p<0.01$, ***$p<0.001$*).

Figure S4

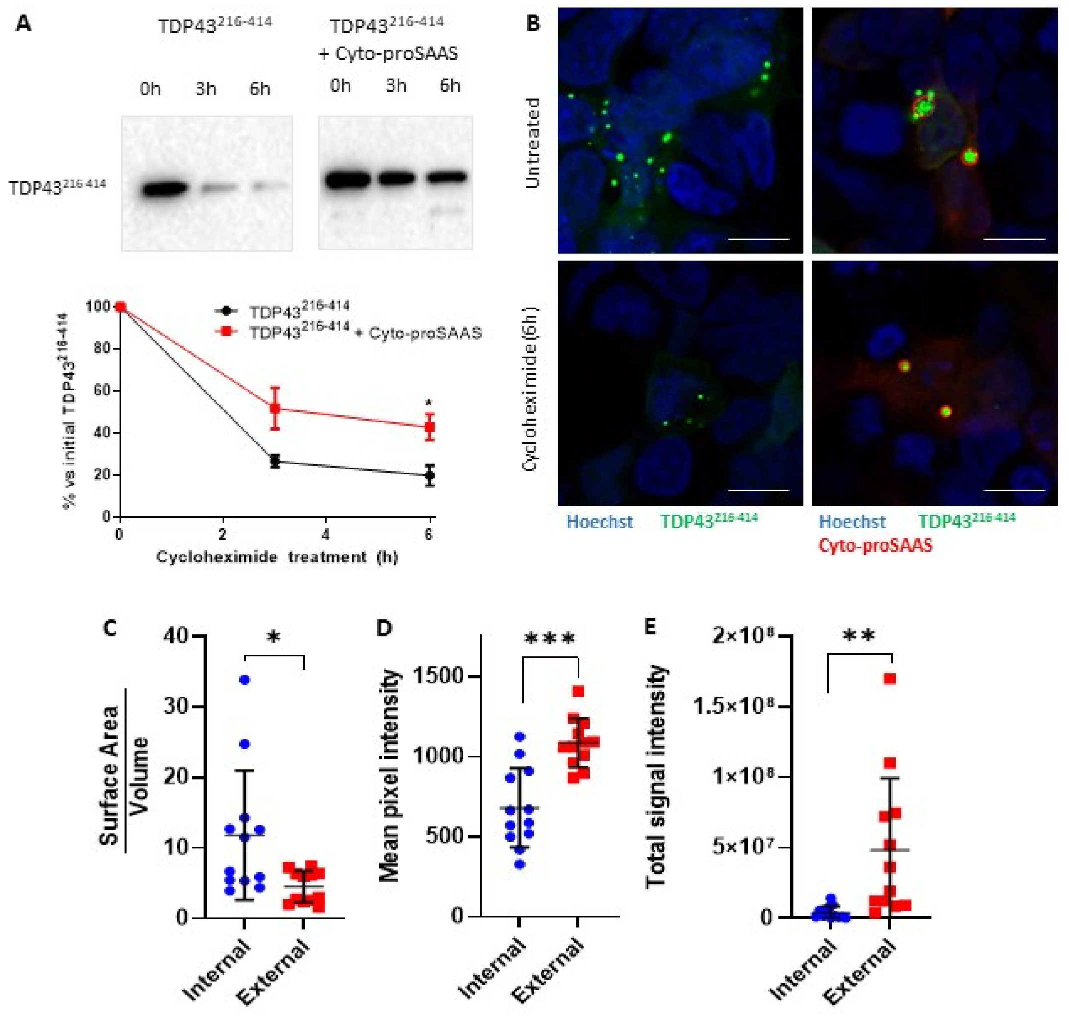

**Supplemental Figure S5: ProSAAS contains intrinsically disordered domains and low complexity regions.**

ProSAAS domains (**Panel A**) containing overlapping low complexity regions (LCRs) (**Panel B**) and intrinsically disordered domains (IDDs) (**Panel C**) as predicted by SEG (PlaToLoCo) and PONDR bioinformatics tools, respectively.

Figure S5

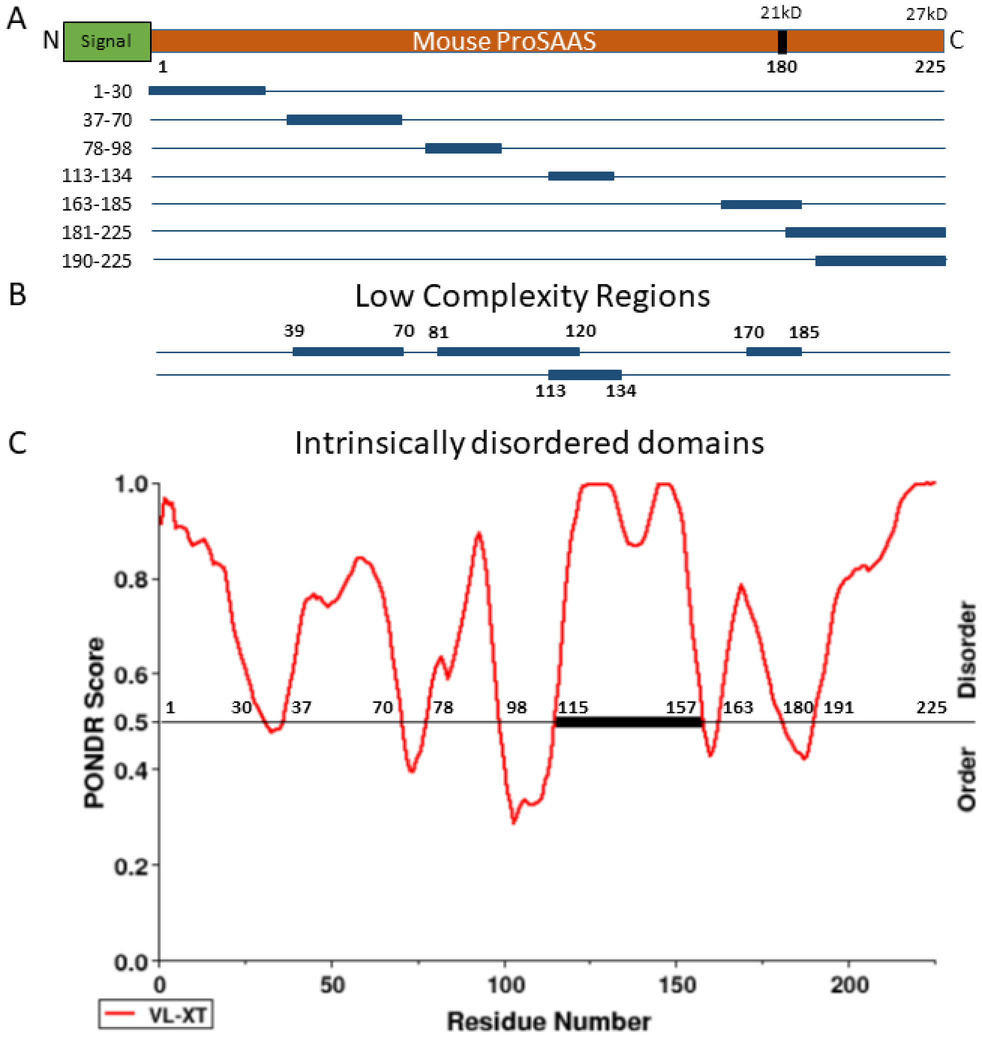
